## Supplementary Information for "The optimal clutch size revisited: separating individual quality from the parental survival costs of reproduction"

**Methods details of brood manipulation data extraction**

Firstly, we re-extracted the raw parental survival values for each given clutch size from the studies in Santos & Nakagawa (2012). This gave a continuous scale of clutch size, as opposed to Santos & Nakagawa (2012) who compared both brood increases and brood reductions, irrespective of the size of these manipulations, as combined categories to the control category of a study. This allowed us to directly compare observational and experimental studies and allowed us to account for the severity of the change in clutch size. We then expanded the number of studies by also including mixed-sex studies, where survival returns were combined for both parents, and included studies published in the years following publication of the Santos & Nakagawa (2012) paper. To do this we used a key word search on Web of Science and Google Scholar using the following terms: “longevity” OR “lifespan” OR “survival” AND "breeding success" OR "brood size" OR "clutch size" OR "number of chicks" OR "number of eggs" AND "trade-off" OR "trade offs" AND fitness AND life-history AND avian OR bird OR birds OR ornithology.

**Details of equations used to calculate selection differentials**

$$S\left( t \right)=e^{-\lambda t}$$

***Equation 1.*** *Simple exponential survival approximation function for when risk is equal, i.e. when annual survival independent of age used. Where S = survival, t = age, lambda (*$\lambda)$ *is annual survival*

Annual survival in the treatment category is changed by increased reproductive effort on a log odd ratio scale. To adjust for these survival consequences we first convert this to a linear scale.

$${OR}_{R}= e^{(ES * R)}$$

***Equation 2.*** *Conversion of logged OR (ES) function of R (reproductive effort, as added chicks) to a linear OR for an added chick. The ES is estimated from meta-analysis and a range is tested to explore parameter space*

$$LRS={\sum_{t=1}^{\infty} {{(R}_{base}+R)* e}^{-\lambda*{OR}_{R} * t}}$$

***Equation 3.*** *Lifetime reproductive success. Combining equation 1 and 2 gives a survival function adjusted for the OR adjusted for reproductive effect (right hand side). When we multiply the adjusted survival function by reproductive effort at each t and sum this we get total LRS. Reproductive effort here is the extra reproductive output (R) together with the focal species brood size* $(R_{base}$ ). *Note we omit the first year when survival is 1, i.e. t=0. This means we assume reproductive effort happens in t=1 and has direct consequences, i.e. individuals die during reproduction and lose their brood.*

**Details of selection effect estimates**

We expressed fitness consequences as a proportional change in lifetime reproductive success (Figure 4). To compare the selection differentials expressed as the standard deviation from the mean, we first calculated the linear difference in fitness per egg. This is relevant as selection has the potential to drive rapid evolution from a phenotypic change of 0.1-0.3 SD in a generation^19^. We assumed a binary population with half producing an extra egg. Note, however, that this probably leads to an overestimate of the fitness effect; the proportion in the population genetically predisposed for this trait is most probably smaller than half. We can then use estimates of the SD to investigate whether rapid selection is expected. Only furthest away from the observed fast–slow pace-of-life continuum (the diagonal of Figure 3) was selection strong enough to come close to this rule of thumb. At an average clutch size of 2, a survival rate of 0.2 and using the meta-analytic effect size observed, -0.05, selection (assuming h^2^ of 1, thus probably overestimating the change in one generation) would result in a 0.09 SD change in the phenotypic mean over one generation. For this example, the species average clutch size would rapidly evolve to be larger. Over several generations, the selection effect would reduce to the point where the species would lie where an extant species is observable - a species with a survival rate of 0.2 and a species average clutch size of 10. At this point, the selection effect reduces to 0.007 SD change from the phenotypic mean over one generation. This selection effect is small and therefore, at a level where environmental and genetic effects could counterbalance (or even reverse periodically) any selection pressures, maintaining the constraint of clutch size to the within-species variation.

Table S1: Model outputs for meta-analyses estimating the effect size of the odds of survival for increasing clutch size given the species average clutch size. Treatment was coded as a categorical variable indicating whether studies were either experimental or observational. The species average clutch size was centred to the average clutch size of all species used in the meta-analysis. The increase in clutch size was modelled as the raw increase in clutch size and the standardised increase in clutch size. We have not presented the proportional increase in clutch size as this represents the change from the species average and so is null in this model.

| Model | Parameter | Effect size | 95% CI  lower bounds | 95% CI  upper bounds | p-value |
| --- | --- | --- | --- | --- | --- |
| Raw | Intercept | -0.047 | -0.147 | 0.054 | 0.363 |
|  | Treatment: Observational | 0.150 | 0.079 | 0.222 | **<0.0001** |
|  | Centred species clutch size | 0.011 | -0.012 | 0.034 | 0.337 |
|  | Treatment: Observational x Species clutch size | -0.036 | -0.066 | -0.007 | **0.015** |
| Standardised | Intercept | -0.065 | -0.222 | 0.092 | 0.418 |
|  | Treatment: Observational | 0.202 | 0.074 | 0.330 | **0.002** |
|  | Species clutch size | 0.015 | -0.026 | 0.055 | 0.482 |
|  | Treatment: Observational x Species clutch size | -0.057 | -0.117 | 0.002 | 0.057 |
| *Model = ~observational_or_experimental * mean_adjusted_clutchsize, random = (species, phylogeny, study reference)* | | | | | |

Table S2: Model outputs for survival given increasing clutch size for brood manipulation (n= 30 (female), 20 (male) and 8 (mixed sex)) and observational studies for the different sexes (n= 11 (female), 7 (male) and 2 (mixed sex)). Mixed sex studies were found to be at the extremes of the trend, a reflection of species who lay smaller clutch sizes rather than an effect of the mixed sex itself.

| Clutch size measure |  | Sex | Estimate | SE | P-value | CI.lb | CI.ub |
| --- | --- | --- | --- | --- | --- | --- | --- |
| Raw | Brood manipulation | Female | -0.0361 | 0.0407 | 0.3754 | -0.1158 | 0.0437 |
|  |  | Male | 0.0186 | 0.0452 | 0.6807 | -0.07 | 0.1072 |
|  |  | Mixed | -0.2079 | 0.0617 | **0.0007** | -0.3287 | -0.087 |
|  | Observational | Female | 0.1279 | 0.104 | 0.2189 | -0.076 | 0.3317 |
|  |  | Male | 0.0232 | 0.1067 | 0.8282 | -0.186 | 0.2324 |
|  |  | Mixed | 0.49 | 0.2638 | 0.0632 | -0.027 | 1.007 |
| Standardised | Brood manipulation | Female | -0.082 | 0.0665 | 0.2179 | -0.2124 | 0.0484 |
|  |  | Male | 0.0318 | 0.0764 | 0.6778 | -0.1181 | 0.1816 |
|  |  | Mixed | -0.2686 | 0.1264 | **0.0336** | -0.5163 | -0.0209 |
|  | Observational | Female | 0.1732 | 0.1491 | 0.2452 | -0.119 | 0.4654 |
|  |  | Male | 0.0076 | 0.156 | 0.9614 | -0.2982 | 0.3133 |
|  |  | Mixed | 0.5216 | 0.3214 | 0.1047 | -0.1084 | 1.1516 |
| Mean adjusted | Brood manipulation | Female | -0.2776 | 0.1909 | 0.1458 | -0.6517 | 0.0965 |
|  |  | Male | 0.1135 | 0.2368 | 0.6318 | -0.3506 | 0.5776 |
|  |  | Mixed | -0.6169 | 0.2773 | **0.0261** | -1.1604 | -0.0734 |
|  | Observational | Female | 0.5721 | 0.3148 | 0.0692 | -0.0449 | 1.1892 |
|  |  | Male | 0.0639 | 0.3287 | 0.8459 | -0.5803 | 0.7081 |
|  |  | Mixed | 0.9363 | 0.5614 | 0.0953 | -0.164 | 2.0366 |
| *Model = observational_or_experimental + sex,* *random = (species, phylogeny, study reference)* | | | | | | | |

Table S3: I^2 values for each model showing the proportion of variation accounted for by the random effects of the model. The phylogenetic signal was included as a correlation matrix within the model.

| Model | I^2 | | | | |
| --- | --- | --- | --- | --- | --- |
|  | Total | Species | Phylogenetic | Reference | Total species effect  (Species + Phylogenetic) |
| Raw | 0.494 | 0.000000003 | 0.287 | 0.208 | 0.287 |
| Standardised | 0.542 | 0.080 | 0.00000002 | 0.463 | 0.080 |
| Proportional | 0.428 | 0.137 | 0.00000001 | 0.291 | 0.137 |

Table S4: Excluded studies and the rationale for exclusion.

| **Reference** | **Species** | **Reason for exclusion** |
| --- | --- | --- |
| Ashcroft, 1979 | *Puffinus puffinus* | No parental survival values given clutch/brood size. Also no clutch/brood size variation in focal species |
| Erikstad et al, 2009 | *Puffinus puffinus* | No clutch/brood size manipulation. Manipulation is age of offspring. |
| Wernham & Bryant, 1998 | *Puffinus puffinus* | No clutch/brood size variation in study |
| Wiebe, 2005 | *Colaptes auratus* | Mate removal, not clutch/brood manipulation |
| Askenmo, 1979 | *Ficedula hypoleuca* | Doesn't state manipulation size |
| Tinbergen & Both, 1999 | *Parus major* | Manipulation is to equalise brood size throughout population |
| Annett & Pierotti, 1999 | *Larus occidentalis* | Breeding lifespan not survival |
| Murphy 2007 | *Tyrannus tyrannus* | No survival values given |
| Lessells, 1986 | *Branta canadensis* | No parental survival values given clutch/brood size. |
| Schaub & Hirschheydt, 2009 | *Hirundo rustica* | Clutch sizes are pooled (0 offspring,1-6 offspring and 6+ offspring) with large variation in each group meaning it is not informative for our study to gain reasonably accurate survival given clutch size raised. |
| Milonoff & Paananen, 1993 | *Bucephala clangula* | Clutch size before manipulation varies significantly |
| Blondel et al, 1998 | *Parus caeruleus* | No parental survival values given clutch/brood size. |
| Knowles, Wood & Sheldon, 2010 | *Parus caeruleus* | No parental survival values given clutch/brood size. |
| Kluyver, 1970 | *Parus major* | Combined first and second broods |

Figure S1: A) The percentage change in brood size of the maximum manipulation (regardless of whether the manipulation was an increase or reduction) in experimental studies given the species average brood size. B) The brood size of each experimental study. Points indicate the species average brood size (i.e., before manipulation occurred), shaded bars indicate the natural standard deviation in clutch size observed in the species (i.e., without manipulation) and the whiskers represent the lower and upper clutch sizes after a manipulation occurred.

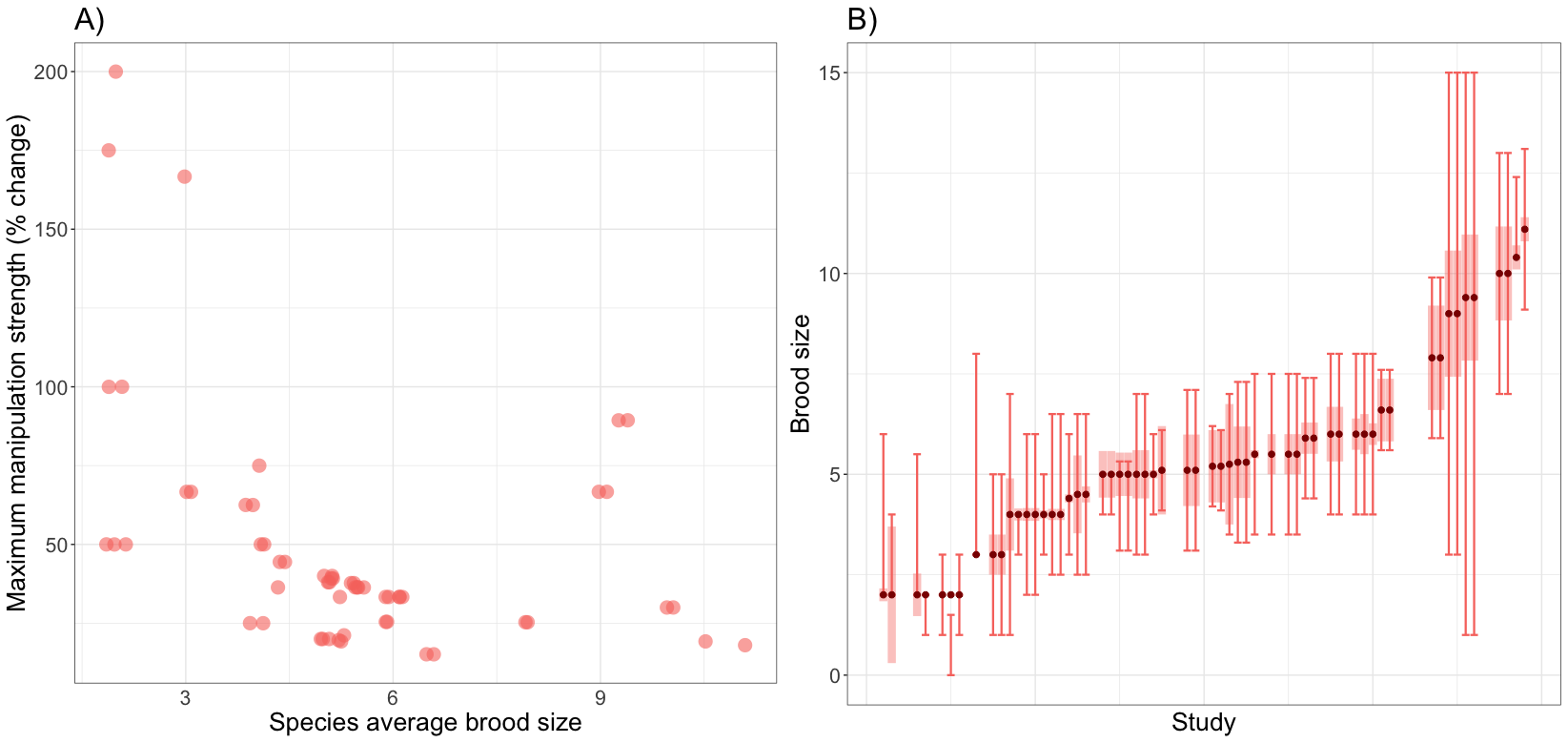

Figure S2: Survival effects for increasing clutch size for female, male and mixed sex studies. The clutch size was measured in three ways; raw clutch size, standardised and mean adjusted. Separate meta-analyses were run for observational (n= 11 (female), 7 (male) and 2 (mixed sex)) and brood manipulation (n= 30 (female), 20 (male) and 8 (mixed sex)) studies. Points are the combined effect size and whiskers are the 95% confidence intervals.

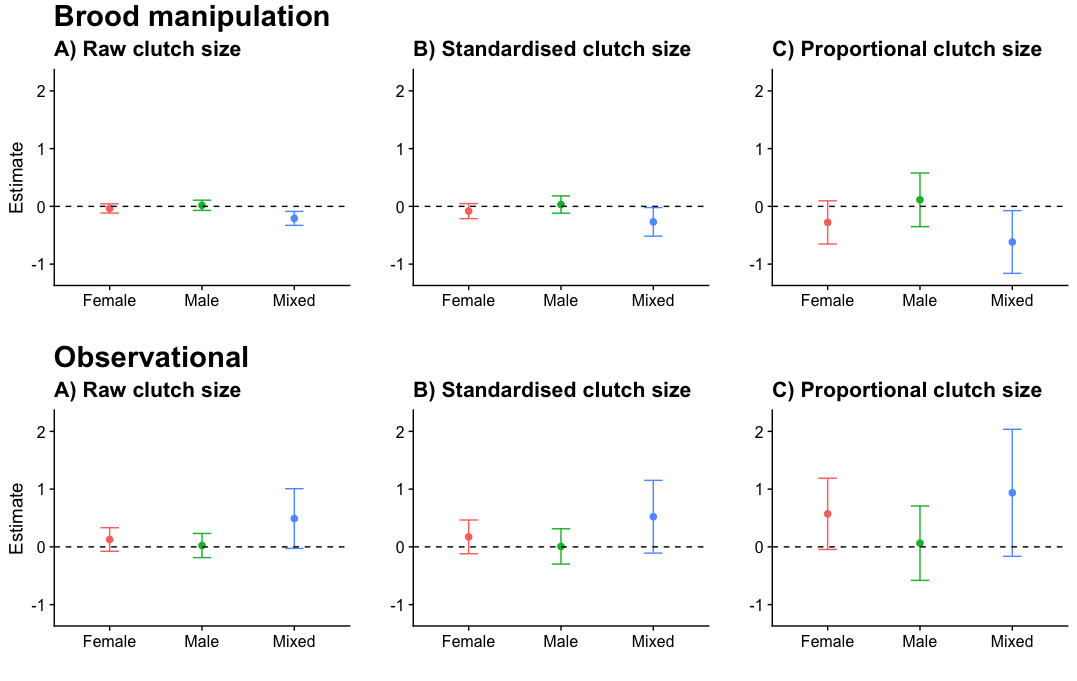

Figure S3: Funnel plot of meta-analysis residuals against standard error. Brood manipulation and observational data are combined. Red points are for experimental studies and blue points are observational studies.

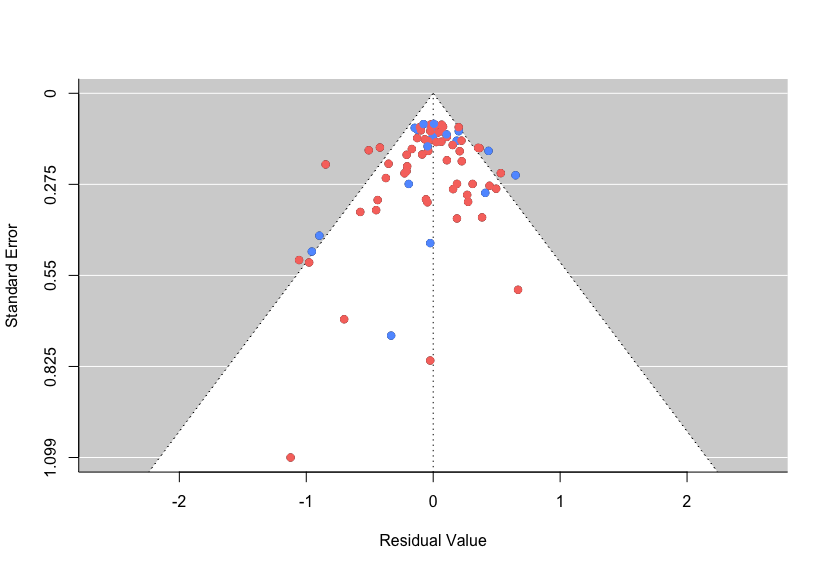
